# Deep Learning Detection of Subtle Torsional Eye Movements: Preliminary Results

**DOI:** 10.1101/2024.05.26.595236

**Authors:** Krishna Mukunda, Tianyi Ye, Yi Luo, Asimina Zoitou, Kyungmin (Esther) Kwon, Richa Singh, JiWon Woo, Nikita Sivakumar, Joseph L. Greenstein, Casey Overby Taylor, Amir Kheradmand, Kemar Earl Green

**Author notes:** Co-first author (Krishna Mukunda, Tianyi Ye).

## Abstract

The control of torsional eye position is a key component of ocular motor function. Ocular torsion can be affected by pathologies that involve ocular motor pathways, spanning from the vestibular labyrinth of the inner ears to various regions of the brainstem and cerebellum. Timely and accurate diagnosis enables efficient interventions and management of each case which are crucial for patients with dizziness, vertical double vision, or imbalance. Such detailed evaluation of eye movements may not be possible in all frontline clinical settings, particularly for detecting torsional abnormalities. These abnormalities are often more challenging to identify at the bedside compared to horizontal or vertical eye movements. To address these challenges, we used a dataset of torsional eye movements recorded with video-oculography (VOG) to develop deep learning models for detecting ocular torsion. Our models achieve 0.9308 AUROC and 86.79 % accuracy, leveraging ocular features particularly pertinent to tracking torsional eye position.

## 1 INTRODUCTION

A complete understanding of how the brain generates eye movements requires measurement of not just horizontal and vertical eye movements, but also the torsional position of the eyes. The torsional eye position or ocular torsion is the orientation of the eye around the anterior-posterior axis (or the line of sight) [1]. Torsional eye movements are particularly important for evaluation of the vestibulo-ocular function in response to head movements against the pull of gravity (i.e., head tilt in the roll plane). During head tilt, dynamic changes in ocular torsion is characterized as torsional nystagmus with slow and fast changes in the torsional eye position (Figure 1). With sustained head tilt, there is a static change in the torsional eye position that represents how the vestibular system can detect and generate an ocular response to the pull of gravity [2][3]. For example, during a left head tilt, the dynamic torsional response consists of a series of slow-phase changes of ocular torsion to the right side, each followed by a fast-phase correction to the left side. With the sustained head to the left side, however, there is only a static torsion to the right side without any fast phase correction. These responses are mainly driven by the vestibular inputs through projections from the labyrinths to the ocular motor centers within the brainstem that generate eye movements [1]. Therefore, the detection and measurement of ocular torsion can be particularly valuable for evaluation of vestibular and ocular motor functions in clinical disorders [4]. Such pathologies may involve the vestibular labyrinth of the inner ears or various regions within the brain stem and cerebellum [5].

**Figure 1.**
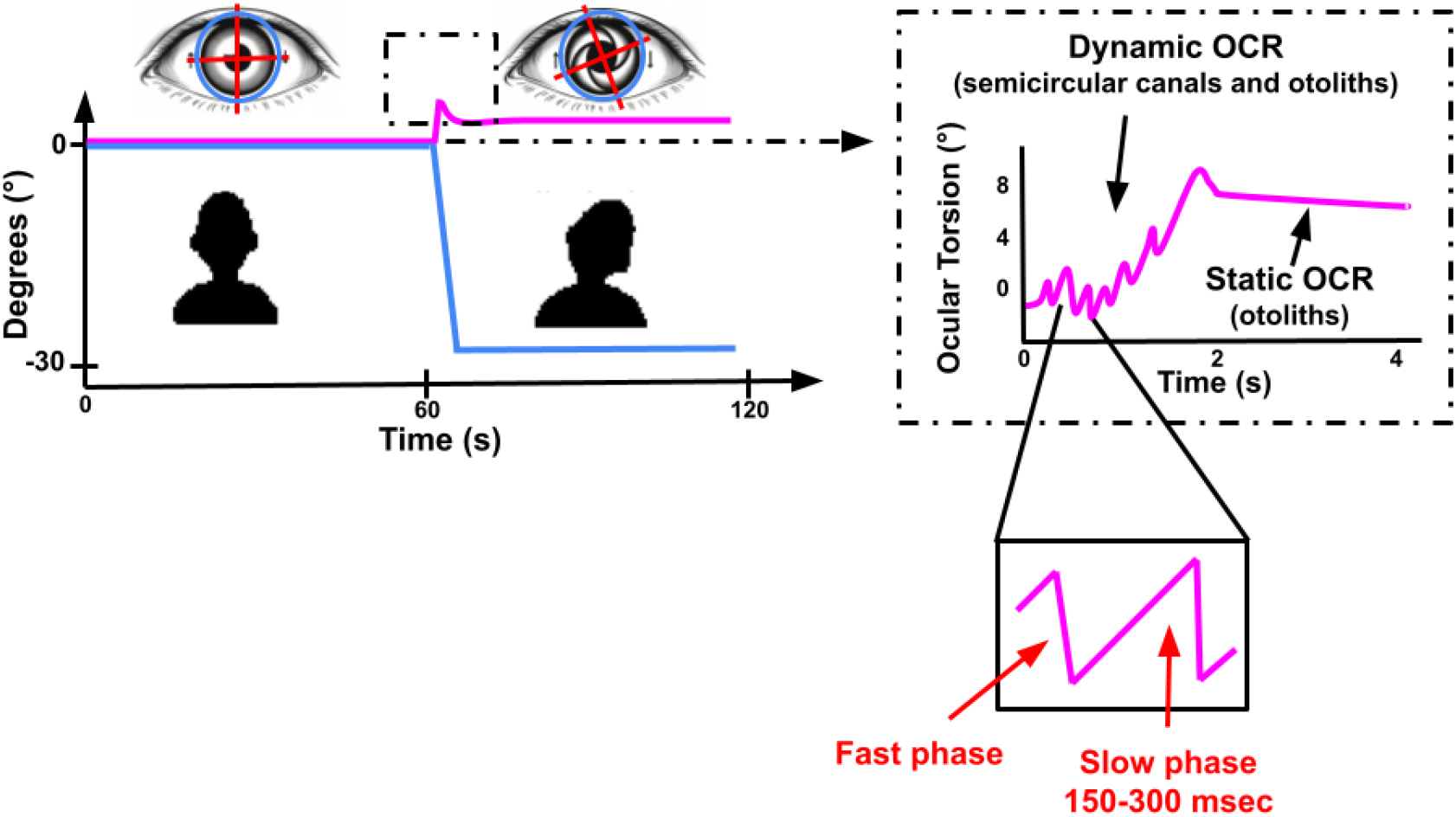
Changes in ocular torsion (magenta) with head tilt (blue) is known as ocular counter-roll (OCR) and consist of dynamic changes during the head tilt and a static change when the head stops moving [8]. The dynamic OCR consist of changes in ocular torsion with slow phases away and fast phases in the direction of the head tilt (i.e., torsional nystagmus). The static OCR consists of a static change in ocular torsion away from the head tilt.

Conventionally, changes in ocular torsion is measured using video ocular counter roll (vOCR) test, which is used for assessment of vestibular function during head tilt [6, 7, 8]. This method relies on tracking iris, which means that occlusion of the iris and pupil by the eyelids must be accounted for to ensure precise measurement of ocular torsion. [9, 10]. While this method achieves high accuracy in clinical settings, the required technology is not widely available in home environments and necessitates trained technicians for administration, thereby hindering rapid testing. Furthermore, other factors such as changes in pupil size, eyelid twitching, and conjunctival vessels may also serve as biomarkers for subtle torsion detection, which have not been accounted for in previous methodologies.

Here, we developed preliminary deep learning models to detect changes in ocular torsion from eye movement recordings. Understanding the predictions from these models can facilitate future detection of ocular torsion by considering a broader range of features beyond those typically used by traditional methods. This approach holds significant value in measuring ocular torsion during the clinical assessment of patients experiencing dizziness or double vision in frontline healthcare settings.

## 2 METHODOLOGY

### 2.1 Data processing and training

Our dataset includes raw binocular video-oculography (VOG) recordings (frame rate = 100Hz; resolution=260 x 400 pixels) and numerical time series data changes in eye position during lateral head tilts. The preliminary datasets consists of 60 videos (12 head movements per video) from 15 subjects from 2 trials of lateral head tilt in each direction. Each video was divided into 500 milliseconds (0.5s) clips. Torsional waveform data consists of multiple slow and fast-phase combinations, known as nystagmus beats, with the duration of a slow-phase being approximately 150-300 milliseconds (Figure 1). A minimum of at least 2 consecutive beats is needed to detect meaningful dynamic torsional movements. Therefore 500 millisecond was chosen as the minimum clip length to capture 2 beats. The class label for each clip was validated using ground-truth from the gold-standard torsional detection method [11].

The dataset construction currently employs an over-down-sampling approach. Oversampling is achieved by using a sliding window of 5 frames when dividing each video into clips, which increases the number of clips per video tenfold (as illustrated in Figure 2.B.). Additionally, the number of clips containing torsional nystagmus can differ as the the duration of duration of the nystagmus may vary among individuals. To ensure an equal number of torsional clips were selected from each video, the video with the fewest clips of torsional nystagmus was identified. Subsequently, all videos were down-sampled to match this minimum number of torsional clips (n = 60 clips per video), thus maintaining a balanced dataset where each video was equally represented in the training of the model. Furthermore, an equal number of clips without changes in ocular torsion were randomly chosen from each video to match the number of torsional clips.The constructed dataset, comprising 7200 clips, was then grouped by subjects to prevent data leakage—where training data improperly includes test information—ensuring that each subject’s four videos recordings (2 for each eye and 2 for each trial) were used exclusively in either the training or testing sets. The dataset was split into different sets for training, validation, and testing that included 5280, 960, and 960 clips respectively. The training set was used to train the model, the validation set was employed to assess the performance of the model during training, and the test set was reserved for final evaluation, not used at any point during the training process.

**Figure 2.**
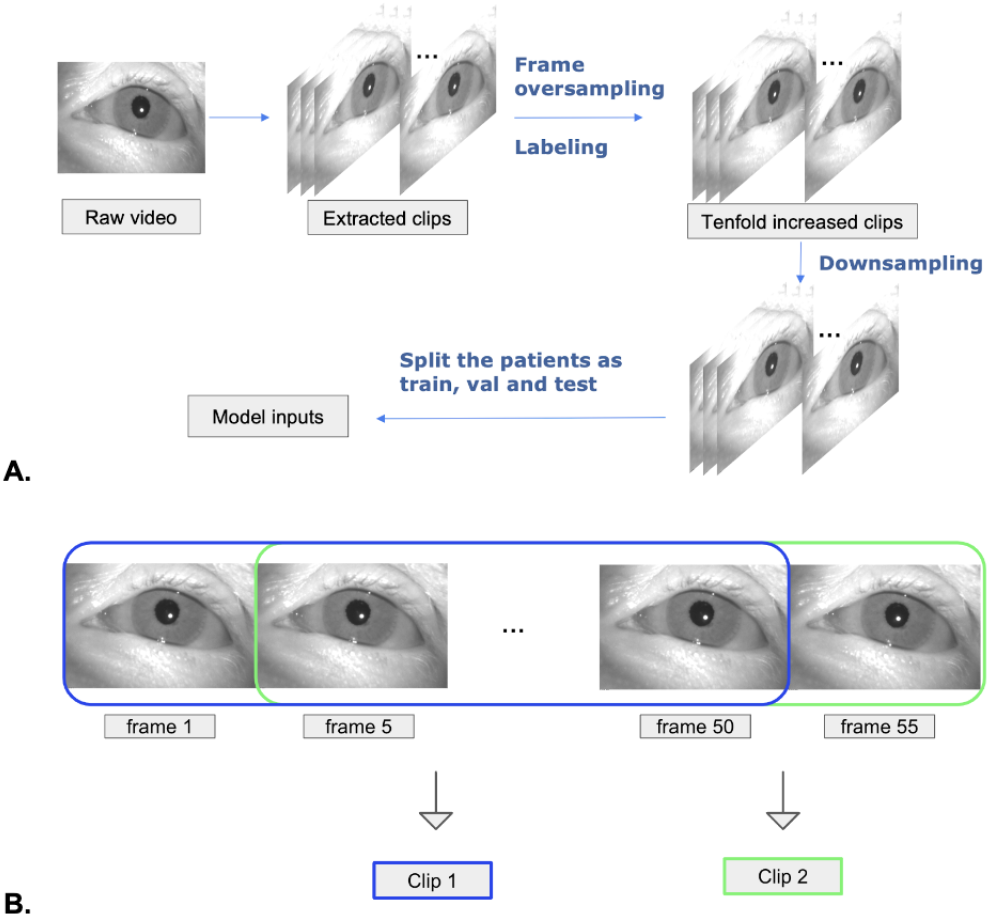
(A) Dataset construction workflow. The raw video is divided into clips using a sliding window technique for oversampling, which increases the number of clips by allowing overlap during extraction. Clips are then downsampled to create a balanced dataset, ensuring equal representation of torsional and non-torsional clips in each video by matching the smallest number of torsional clips found across all videos. The dataset is subdivided into training, validation and testing sets for model training and evaluation. (B) Overview of frame oversampling method using a sliding window. The number of clips is over sampled so that every 5 frames a new 50 frame window is generated, denoting a clip. This allows for a 10 fold increase in the number of clips.

### 2.2 Development and evaluation of deep learning models

The ResNet-18 convolutional neural network (Fig. 3C) was used as the baseline model for further training since it is a generally good architecture for binary classification tasks [10]. In this study, we used 2D and 2.5D Residual Network (ResNet) models for training. The 2.5D approach inputs a temporal sequence of frames into a 2D ResNet architecture, effectively embedding temporal information into the spatial domain early in the model training. This allows the 2D Convolutional Neural Network (CNN) to process temporal data efficiently. This approach allows the network to consider temporal information without fully committing to a 3D convolutional structure, which processes data in three dimensions, accounting for spatial and temporal features simultaneously. It offers a compromise between computational efficiency and the ability to leverage temporal dynamics.

**Figure 3.**
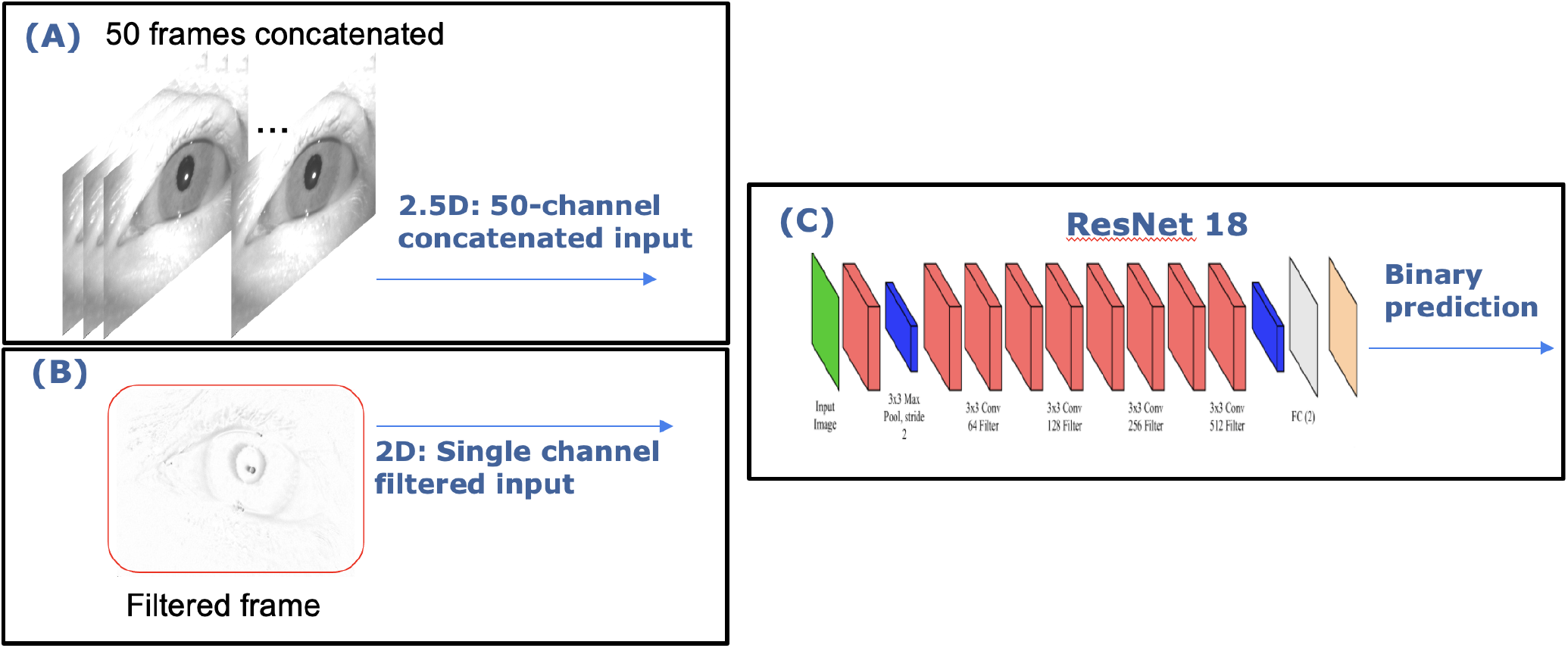
(A) Input of 2.5D ResNet, (B) Input of 2D ResNet and (C) architecture of ResNet 18. The input of the 2.5D ResNet model was the full 50 video frame clip. The input of the 2D ResNet model was based on the recursive filtering method to generate one filtered image encompassing all the motion from 50 video frames.

#### 2.2.1 2.5D ResNet based on consecutive frames

One approach for achieving a binary classifier for torsion detection involves the development of a 2.5D ResNet model. The 2.5D approach aims to bridge the gap between 2D and 3D analysis by using a sequence of frames as input to a 2D ResNet architecture. While 3D analysis incorporates a temporal dimension throughout the model architecture, the 2.5D model integrates the temporal information of sequential frames into the spatial domain early in the processing. This results in temporal processing occurring early in the model training, as opposed to 3D models that incorporate temporal information throughout the training process. Here we used 50 consecutive video frames concatenated along the temporal dimension to form a 3D input, thereby encapsulating the ocular torsion pattern in the video clip. In this approach, the model can detect ocular torsion based on both morphological and temporal features. To enhance the diversity of the data and prevent the model from memorizing specific patterns, we introduced two augmentation techniques: random horizontal flip and rotation by 0-5 degrees. The random horizontal flip mirrors the images along the vertical axis, simulating different viewing angles. The rotation by 0-5 degrees slightly tilts the images, mimicking natural variations in camera or subject positioning. These augmentations help the model generalize better to unseen data by exposing it to a wider range of possible scenarios. We used batch size (the number of clips shown to the model at once during training) = 64 and learning rate (the magnitude of updates made to the model’s weights during training) = 0.0001. When the Area Under the Curve (AUC, a metric evaluating model performance) on the validation set did not increase for 2 epochs (complete passes through the training dataset), we halved the learning rate. If there was no increase for 20 epochs, we stopped the training. The best model was selected based on the highest AUC value on the validation set.

#### 2.2.2 2D ResNet based on the recursive filtering method

Here we segmented videos into 50-frame clips to construct filtered images. The filtered image highlights movements within a video clip by identifying differences in each frame recursively [12, 13]. This means that in each clip, pixel values are cumulatively aggregated up to the current frame, *M*_*t*_, at time *t*, forming an intermediate frame, *I*_*t*_. The current frame is then compared to this intermediate frame to calculate pixel differences. These differences are subsequently amplified by a beta value, *β*, that enhances the distinction between frames. The cumulative result of these amplified differences constitutes the filtered image, *F*_*t*_. The equation for the calculation of the filtered image is as follows:

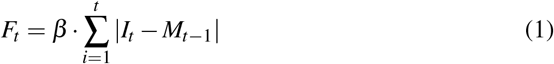

where |*I*_*t*_ − *M*_*t*−1_| represents the absolute difference between the current frame and the intermediate frame at time *t*, and *β* is the amplification factor.

These filtered images are then analyzed by a two-dimensional ResNet model, which processes the data to generate binary predictions that discern the presence of torsional movement within each frame. This approach was adopted from a method by Wagle et al method for detection of nystagmus [14].

## 3 RESULT

### 3.1 Model performance

Our 2.5D model shows a torsion and non-torsion clip differentiation accuracy=0.8679, AUC=0.9308, sensitivity=0.8962, and specificity=0.8396.

The 2D model has a torsion and non-torsion clip differentiation accuracy=0.81, AUC = 0.88, sensitivity=0.84, and specificity=0.78.

### 3.2 Model interpretability

The interpretability of the models is a critical element that pertains to the features used for the detection of torsion and allows confidence in their decision-making processes. To evaluate interpretability, we utilized Gradient-weighted Class Activation Mapping (Grad-CAM) [15] of the final convolutional layer of each model for the task of torsion detection. As Figure 4 and Figure 5 show respectively for the 2D and 2.5D models, Grad-CAM generates a heatmap that visualizes the significance of each pixel in an image. These heatmaps are determined by the model’s weights to illustrate the specific regions that the model focuses on during its predictive analysis. The heatmap values are measured as normalized activation intensity, indicating the relative importance of different regions in the image for the model’s decision, normalized to a scale from 0 to 1 [15]. Both the 2D and the 2.5D trained ResNet models appear to focus primarily on the iris features. Physiological torsion in a clinical setting is primarily identified though the observation of the iris. Thus, these models appear to use clinically relevant features in their predictive analysis.

**Figure 4.**
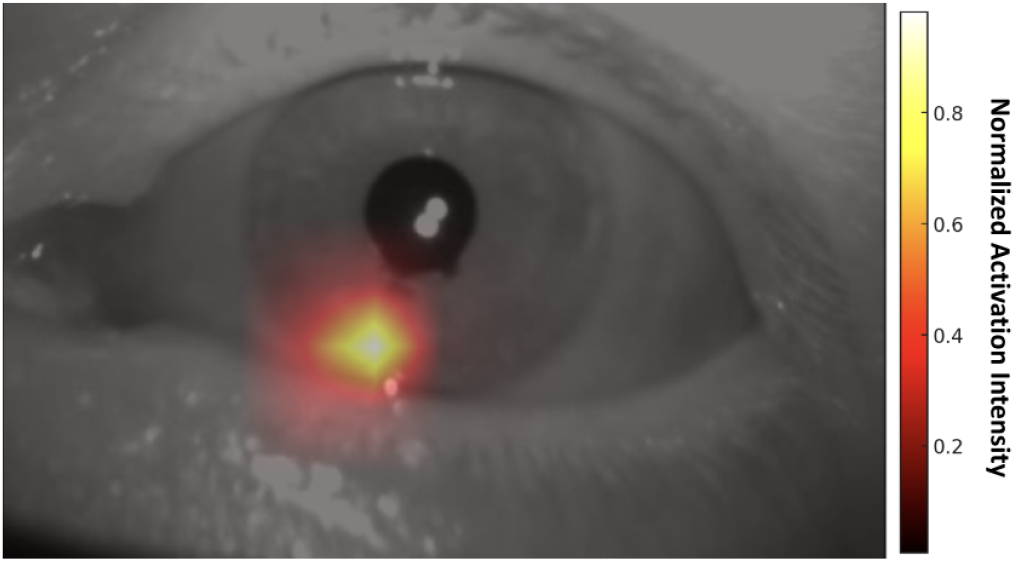
Heatmap generated by Grad-CAM for 2.5D ResNet model showing “segmental” iris focus.

**Figure 5.**
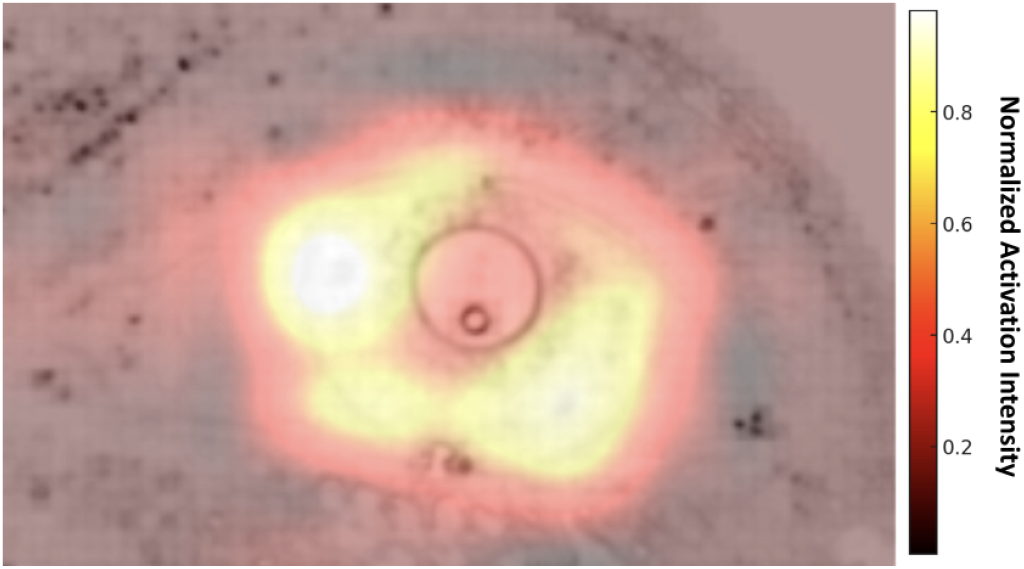
Heatmap generated by Grad-CAM for 2D ResNet showing quasi-circumferential iris focus.

## 4 DISCUSSIONS

Advanced deep learning techniques and interpretable methodologies were employed for the accurate detection of torsional nystagmus. Training of the 2D and 2.5D ResNet on VOG recordings of dynamic torsion from vOCR yielded an accuracy in torsional detection of 0.8679 and 0.81 with AUCs of 0.9308 and 0.88 respectively. Interpretitability of the these methods using Grad-CAM revealed that they focused primarily on the iris, a clinically relevant feature in torsional nystagmus diagnosis. The 2D model applies a recursive filter to a sequence of images to detect changes or movements between frames, effectively generating a “filtered” image that highlights areas of eye motion [14]. This method is computationally more efficient than the 2.5D approach while still being capable of detecting motion. However, this method may not capture the full complexity of ocular torsion, especially in cases where torsion is minimal or occurs alongside other eye movements that might confuse the filter. The incorporation of motion information can lead to improved detection of ocular torsion by learning subtle and complex torsion features using the CNN. However, while the temporal information is incorporated to the 3D input by this method, the temporal dynamics are not fully captured as in a true 3D way - the temporal information is still processed along with spatial information in 2D CNN. In the future, we aim to apply a 3D ResNet model with 3D convolutional layers to disentangle the temporal information from spatial information, therefore enabling a more accurate analysis of dynamic patterns over time, which could lead to significantly improved detection and understanding of diverse patterns of changes in ocular torsion.

